# Iron and zinc biofortification in potato through the introduction of *NICOTIANAMINE SYNTHASE* and *FERRITIN* genes

**DOI:** 10.1101/2025.03.01.640813

**Authors:** Juan I. Cortelezzi, Martina Zubillaga, Victoria R. Scardino, Marina Fumagalli, María N. Muñiz García, Daniela A. Capiati

## Abstract

Micronutrient deficiencies, such as those involving iron, zinc, and vitamin A, represent a critical global public health challenge. Crop biofortification offers a cost-effective solution by enhancing the micronutrient content of staple foods, particularly in regions with limited access to diverse diets. In this study, we developed a high-iron and high-zinc Spunta potato variety through the co-expression of the *Arabidopsis thaliana NICOTIANAMINE SYNTHASE 1* (*AtNAS1*) gene under a constitutive promoter and the *Phaseolus vulgaris FERRITIN* (*PvFERRITIN*) gene under a tuber-specific promoter. Of the five co-expressing events evaluated, we identified FN4 as the optimal line, achieving significant levels of iron and zinc biofortification without compromising yield. This line exhibited an enhanced capacity to accumulate iron in tubers under conditions of increased soil iron availability, maintaining yield even when vegetative growth was reduced due to elevated soil iron levels. The FN4 variety showed a 2.1-and 1.8-fold increase in iron and zinc content, respectively, in the tubers. A typical portion of this variety could provide 33.3% of the recommended dietary allowance (RDA) for iron and 19.8% for zinc in women of reproductive age. This biofortified potato has the potential to reduce iron deficiency anemia and zinc deficiency, enhancing the health of populations in both rural and urban areas affected by poverty.

## INTRODUCTION

Deficiencies in essential micronutrients, such as iron, zinc, vitamin A, folate, vitamin B12, vitamin D, and iodine, can lead to severe health consequences (Stevens et al. 2022). According to the World Health Organization (WHO), iron deficiency anemia is the most common nutritional disorder, affecting approximately 25% of the global population, with young children and pregnant women at greatest risk (WHO 2015). Zinc deficiency is another major concern, particularly as it contributes to stunting in children (International Zinc Nutrition Consultative Group 2004). It is estimated that 1.1 billion people worldwide are at risk of zinc deficiency due to inadequate dietary intake (Kumssa et al. 2015).

Crop biofortification has emerged as a promising strategy to tackle food security challenges, particularly in developing countries (Labuschagne 2023). This approach enhances the nutritional value through selective breeding or biotechnological methods to increase essential nutrients in the edible parts of plants. Advances in genetic engineering to enhance iron and zinc levels in crops began in the early 21^st^ century, with various methods tested over time (Stangoulis and Knez 2022). Early strategies for iron biofortification focused on the use of the *FERRITIN* gene, which encodes a storage protein capable of binding up to 4,500 iron atoms (Theil 2003). In rice, *FERRITIN* genes from *Glycine max or Phaseolus vulgaris* were expressed using endosperm-specific promoters, leading to a 2-to 3-fold increase in grain iron concentration (Goto et al. 1999; Lucca et al. 2002; Vasconcelos et al. 2003). However, other studies reported no improvements in rice iron concentration using the same approach (Qu Le et al. 2005). The introduction of *FERRITIN* alone may be insufficient to significantly enhance iron content, as effective biofortification likely requires not only improved iron storage but also increased uptake and translocation. In this context, the *NICOTIANAMINE SYNTHASE* (*NAS*) gene has gained attention. Nicotianamine (NA), a non-proteinogenic amino acid, chelates metal cations and plays a crucial role in the internal transport of iron and zinc within plants (Nozoye 2018). Introducing *NAS* alone has led to iron content increases of 2-to 4.5-fold in soybean seeds, sweet potato tubers, potato tubers, rice grains, and wheat grains (Nozoye et al. 2014; Nozoye et al. 2017; Zha et al. 2022; Masuda et al. 2009; Johnson et al. 2011; Singh et al. 2017). This strategy also provides the additional benefit of increasing zinc levels (Nozoye 2018). In rice, the combined expression of *NAS* and *FERRITIN* has achieved significantly higher iron biofortification compared to *FERRITIN* alone, resulting in a 4-to 6-fold increase in iron content in grains (Tsakirpaloglou et al. 2023; Wirth et al. 2009). However, in wheat, co-expression of *NAS* and *FERRITIN* did not result in higher iron levels compared to the expression of either *NAS* or *FERRITIN* alone (Singh et al. 2017). These findings highlight the need for biofortification strategies to be tailored and tested for each specific crop species.

To further enhance iron biofortification, additional genes have been introduced in combination with *FERRITIN* and *NAS*. One such gene is *OsYSL2*, which encodes a transporter for the Fe(II)-NA complex (Masuda et al. 2012). Another key gene is *IDS3*, which encodes mugineic acid synthase, an enzyme crucial for the biosynthesis of mugineic acid, an iron chelator with functions similar to NA (Masuda et al. 2013). The *AtIRT* gene, encoding the Iron-Regulated Transporter responsible for iron uptake from soil into roots, has also been used in combination with *FERRITIN* and *NAS* (Boonyaves et al. 2017). More recently, the *TaVIT2-D* gene, encoding a vacuolar iron transporter, was introduced alongside *NAS* to enhance iron content in wheat grains (Harrington et al. 2023).

Potato (*Solanum tuberosum*) is the world’s most consumed non-grain food crop, with over 50% of production occurring in developing countries where micronutrient malnutrition is widespread (Singh et al. 2021). Enhancing the micronutrient content of potato tubers is therefore vital for improving public health, particularly in regions where potatoes constitute a dietary staple for a significant portion of the population. While potatoes are already a rich source of micronutrients (Zaheer and Akhtar 2016), they offer a unique advantage in terms of iron absorption. Their iron absorption rate is approximately 28% (Jongstra et al. 2020), substantially higher than the 1-4% typically found in legumes (DellaValle et al. 2015). This makes potatoes a particularly efficient dietary source of iron, which is especially important in areas where iron deficiency is a prevalent public health concern.

This study aimed to develop biofortified varieties of the potato cultivar Spunta, the most widely cultivated variety in Argentina, with enhanced iron and zinc content in its tubers. To achieve this, two genes were employed: the *FERRITIN* gene from *Phaseolus vulgaris* (*PvFERRITIN*), expressed under the control of the tuber-specific *PATATIN* promoter, and the *NAS1* gene from *Arabidopsis thaliana* (*AtNAS1*) regulated by the constitutive CaMV 35S promoter. These genes were introduced either individually or in combination.

## MATERIALS AND METHODS

### Generation of potato transgenic lines expressing *AtNAS1* and *PvFERRITIN*

Construction of the pPZP-NPTII-35S::AtNAS1 vector: The 35S::AtNAS1-tNOS cassette was amplified from the IINF vector (Boonyaves et al. 2017) and cloned into the pPZP-NPTII binary vector.

Construction of the pPZP-NPTII-B33::PvFERRITIN vector: The *PATATIN* promoter (B33) sequence was obtained by restriction enzyme digestion of the pBIN-B33 vector (Rocha-Sosa et al. 1989). The PvFERRITIN-tNOS cassette was amplified by PCR from the IINF vector. The B33::PvFERRITIN-tNOS cassette was assembled in an intermediary vector and then transferred into the pPZP-NPTII binary vector.

Construction of the pPZP-NPTII-AtNAS1-PvFERRITIN vector: The B33::PvFERRITIN-tNOS cassette described above was inserted into the pPZP-NPTII-35S::AtNAS1 vector.

The binary vectors described above were used to generate transgenic plants. Transformation of *Solanum tuberosum* (cv. Spunta) minituber discs was performed using *Agrobacterium tumefaciens*, following the protocol described by Muñiz García et al. (2014). Regenerated plants carrying no plasmid but obtained by the same regeneration method as the transgenic lines were used as controls (referred to as wild type) for phenotypic evaluation. Eight transgenic lines were obtained: one expressing *AtNAS1* (N1), two expressing *PvFERRITIN* (F1 and F2), and five expressing both genes (FN1-FN5).

### Plant growth conditions

Transgenic and wild type plants were propagated *in vitro* from single-node cuttings on MS medium (prod no. M519, PhytoTechnology Laboratories, Shawnee Mission, KS, USA) containing 20 g L^-1^ sucrose solidified with 0.7% (w v^-1^) agar, and cultivated in a growth chamber under a 16-h photoperiod (4,000 lx light intensity) at 22 °C. Soil-grown potato plants were obtained by soil transfer of *in vitro* grown plants (*ex vitro*), or by planting seed tubers. Plants were cultivated in a greenhouse maintained between 22 °C and 24 °C, under a 16-h light photoperiod, in 1 L pots filled with commercial substrate Klassman TS 085 (Klasmann-Deilmann GmbH, Germany).

For the assessment of line FN4 in soil (Fig. 8), plants grown from seed tubers were cultivated in 3 L pots filled with a commercial soil-organic compost mixture (Hi-Soil, Argentina).

To evaluate the effect of soil iron concentration (Fig. 9), plants obtained *ex vitro* were cultivated in 1 L pots containing 200 g of Klassman TS 085 substrate, either with or without the addition of 5 or 10 g of the iron chelate iron(III) ethylenediamine di(o-hydroxyphenylacetic) acid (Fe-EDDHA; Basafer® Plus, COMPO EXPERT, Germany).

### Molecular characterization of transgenic plants

PCR amplification of the transgenes: Plant genomic DNA was isolated from leaves of *in vitro* plants as described by Murray and Thompson (1980). PCR amplification of the transgenes was performed using the primer pairs AtNAS1 (forward and reverse), PvFERRITIN (forward and reverse) and NPTII (forward and reverse). Amplification of elongation factor 1-α gene (*EF1-α*) with the primers EF1-α (forward and reverse) was performed as control. The primer sequences are shown in Supplementary Table S1.

Reverse transcription-PCR (RT–PCR): Expression of the transgenes was determined by RT-PCR. RNA was isolated from leaves and tubers, and cDNA synthesis was performed as described in País et al. (2010). The amount of cDNA used in each reaction was derived from 50 ng of total RNA. PCR reactions were performed using the primer pairs AtNAS1 (forward and reverse), PvFERRITIN (forward and reverse) and EF1-α (forward and reverse) (Supplementary Table S1). Reactions were carried out under the following conditions: *AtNAS1*: 55 °C of annealing temperature, 36 cycles (tubers) - 52 °C of annealing temperature, 28 cycles (leaves); *PvFERRITIN*:

62 °C of annealing temperature, 40 cycles (tubers); *EF1-α*: 50 °C of annealing temperature, 38 cycles (tubers) - 52 °C of annealing temperature, 28 cycles (leaves).

PCR and RT-PCR products were separated by electrophoresis in a 1.5 % agarose gel and visualized by ethidium bromide staining, inverting the original image.

### Growth, morphological parameters and tuber yield

Growth and morphological parameters were determined in soil-grown potato plants obtained either by soil transfer of *in vitro* grown plants (*ex vitro*) or by planting seed tubers. Growth and morphological parameters were determined 4 weeks after soil transfer, or 4 weeks after tuber planting. The apical leaflet area and shape (length to width ratio) of the second fully expanded leaf were determined using the Image J software (Schindelin et al. 2012). Tuber yield was assessed in plants either transferred to soil *ex vitro* or grown from seed tubers. Tubers were harvested after plant senescence to determine tuber yield, evaluate tuber characteristics, and measure metals content.

### Dry matter, protein and starch content

Dry matter content was determined by oven-drying finely chopped tubers at 80 °C until a constant weight was achieved (approximately 16 hours). The dry matter percentage (%) was calculated using the formula: 100 × (dry weight fresh weight^-1^). Protein content in fresh tubers was determined using the Bradford method (Bradford 1976). Starch was measured according to the method described by Muñiz García et al. (2022).

### Determination of iron and zinc content

Iron and zinc content was determined by atomic absorption spectrometry. Approximately 2 g of tuber samples were incinerated at 500 °C for 5 h. The resulting ashes were suspended in 5 mL of 20% (v v^-1^) HCl. The solution was then diluted to a final volume of 50 mL with distilled water. Metal concentrations were measured using an AAAnalyst 200 (Perkin-Elmer) atomic absorption spectrometer. Alternatively iron content was measured using a modified protocol from Manzanares et al. (1990). Tuber samples were ground in liquid nitrogen and extracted with 3 µL of 1N HCl per mg of leaf tissue for 16 h at 20 °C. Following extraction, 300 µL of the solution were incubated with 200 µL of 20% NH₂OH·HCl for 5 min, then buffered with 200 µL of an acetate-acetic acid solution. From this mixture, 200 µL were transferred to a transparent 96-well plate in triplicate. The iron content was determined by adding 100 µL of 1.5% (w/v) o-phenanthroline solution and measuring the absorbance at 510 nm, using a calibration curve generated from iron(III) nitrate processed though the same protocol. This colorimetric method yields results comparable to those of atomic absorption spectrometry in plant samples (Manzanares et al. 1990). The method used for iron measurement is specified in the respective figure legends.

### Measurement of leaf water loss and relative water content (RWC)

Water loss and RWC were determined in the apical leaflet of the second fully expanded leaf from plants obtained *ex vitro* or grown from seed tubers, 4 weeks after soil transfer or 4 weeks after tuber planting. To measure water loss by air drying, leaflets were detached and placed at 22 °C. The fresh weight was measured at different time points. Water loss was calculated as the percentage of initial fresh weight lost at each time point. RWC was determined as described in Muñiz García et al. (2018).

### Stomatal size, stomatal density and stomatal index

Assessments of stomatal characteristics were conducted in plants obtained *ex vitro* or developed from seed tubers, 4 weeks after soil transfer or 4 weeks after tuber planting, respectively. The third fully expanded leaf was collected and impressions of the central area of the abaxial surface were obtained using dental silicone polymer. After drying, these impressions served as molds onto which transparent nail polish was applied. The resulting nail polish film was then transferred to a microscope slide and examined under a Nikon Eclipse Ti-E inverted microscope with 400x magnification. The images acquired were used to quantify stomatal density (number of stomata per mm²), stomatal size (µm²; measured using the Image J software), and the stomatal index, calculated as the number of stomata per 200 x 200 µm area divided by the total number of epidermal cells, multiplied by 100. For each leaf, data from two fields were averaged.

### Chlorophyll and carotenoid content

Chlorophyll (chlorophyll a, chlorophyll b, and total chlorophyll) and total carotenoid content were determined spectrophotometrically. Samples were homogenized in acetone 80% (v v^-1^) (1 mL per 10 mg of fresh weight) followed by centrifugation at 3000 *g* for 5 min. The absorbance was measured at 663, 645 and 480 nm. The content of chlorophyll a, b, and total chlorophyll were calculated according to Arnon (1949). Total carotenoid content (mg g fresh weight^-1^) was calculated using the following formula: [A480 + (0.114 A663) - (0.638 A645)] V (1000 FW)^-1^, where V is the total extract volume (mL), and FW is the weight (mg) of the sample extracted. Alternatively, chlorophyll content was measured on fully expanded leaves using a SPAD chlorophyll meter (Clorofilio1, Cavadevices, Argentina).

### Statistical analysis

Statistical analysis was conducted using one-way ANOVA followed by Bonferronís multiple comparisons test, two-way ANOVA followed by Tukeýs multiple comparisons test, or Student‘s *t* test. A *P-*value < 0.05 was considered statistically significant.

## RESULTS

### Generation of potato transgenic lines expressing *AtNAS1* and *PvFERRITIN*

*S. tuberosum* cv. Spunta was transformed with three distinct plasmids encoding *AtNAS1*, *PvFERRITIN*, or both *PvFERRITIN* and *AtNAS1*, respectively (Fig. 1a). *AtNAS1* was driven by the constitutive 35S promoter from cauliflower mosaic virus, while *PvFERRITIN* was expressed under the control of the tuber-specific *PATATIN* promoter. Eight transgenic lines were successfully established: one expressing *AtNAS1* (N1), two expressing *PvFERRITIN* (F1 and F2), and five co-expressing both genes (FN1-FN5). All transgenic lines exhibited root development in selective media (Supplementary Fig. S1). The presence of the transgenes was confirmed by PCR amplification from genomic DNA (Fig. 1b). Expression of the respective transgenes was confirmed by RT-PCR in tubers and leaves (Fig. 1c, d).

**Fig. 1.**
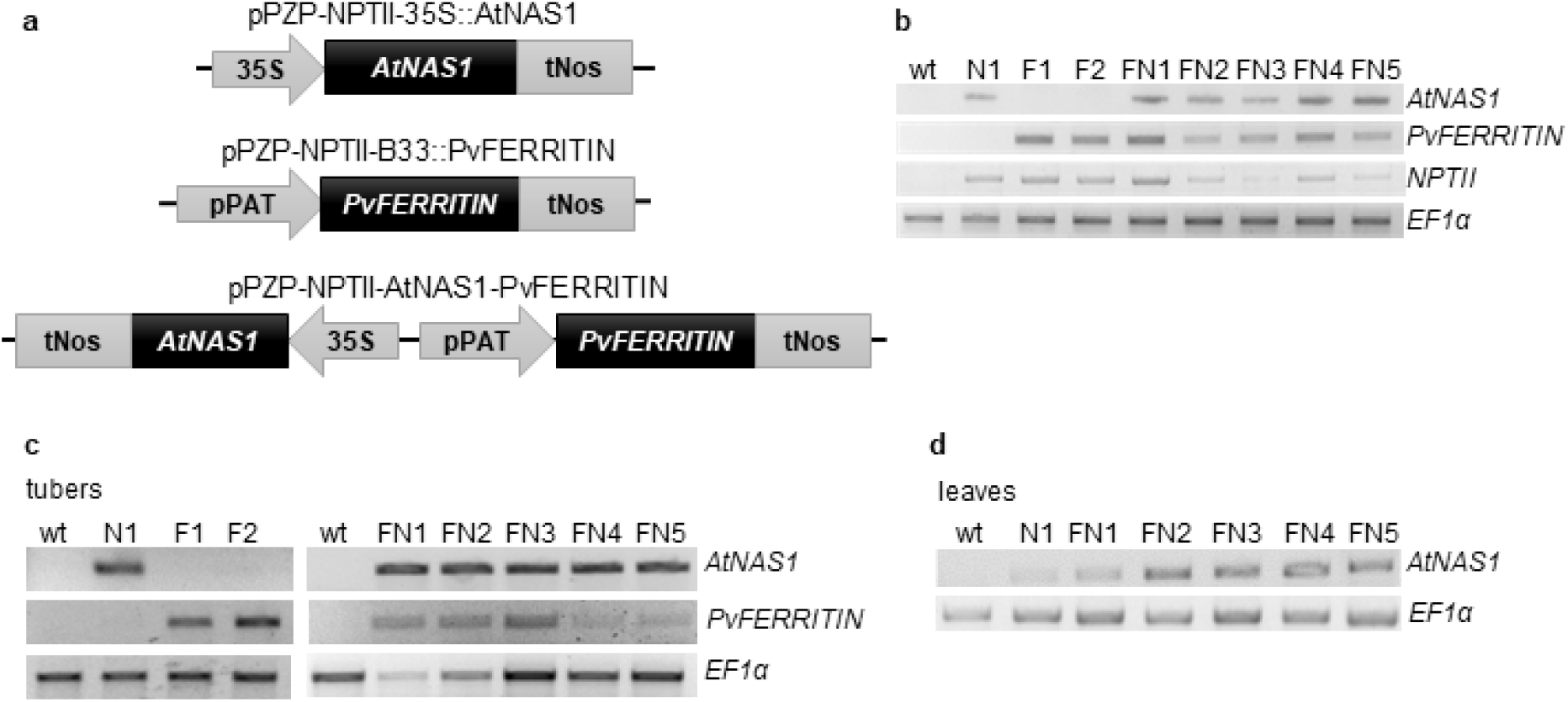
Molecular characterization of the transgenic plants. **a** Schematic representation of the gene constructs used for the development of transgenic plants. **b** PCR analysis of genomic DNA isolated from leaves of wild type (wt) and transgenic plants (N1, F1-2, FN1-5) to detect the presence of the transgenes *AtNAS1*, *PvFERRITIN*, and the *NPTII* gene. **c** and **d** RT-PCR analysis of *AtNAS1* and *PvFERRITIN* in tubers and leaves of wild type and transgenic lines. A representative result of three independent experiments is shown. In **d**, all lanes were from the same gels but were not originally adjacent.

### Vegetative growth and tuberization

The impact of *AtNAS1* and *PvFERRITIN* expression on vegetative parameters was evaluated in plants transferred from *in vitro* conditions to soil (*ex vitro*). Overall, the co-expressing lines exhibited altered growth parameters compared to the wild type (Fig. 2). Lines FN2, FN3, and FN5 displayed reduced stature (Fig. 2a), while FN1, FN3, and FN5 had fewer leaves per plant (Fig. 2b). Additionally, leaf size was smaller in FN1, FN2, FN3, and FN5 compared to the wild type (Fig. 2c, d). Notably, these parameters were unaffected in line FN4 (Fig. 2a-d). Despite the reduction in leaf size in certain transgenic lines, overall leaf morphology remained comparable to that of the wild type across all lines (Fig. 2e, f). A similar trend was observed in plants derived from seed tubers (Supplementary Fig. S2). However, under these conditions, only FN2 exhibited reductions in both leaf number and leaf size (Supplementary Fig. S2b, c), and alterations in leaf morphology were observed in FN1, FN2, FN3, and FN5 (Supplementary Fig. S2d, e).

**Fig. 2.**
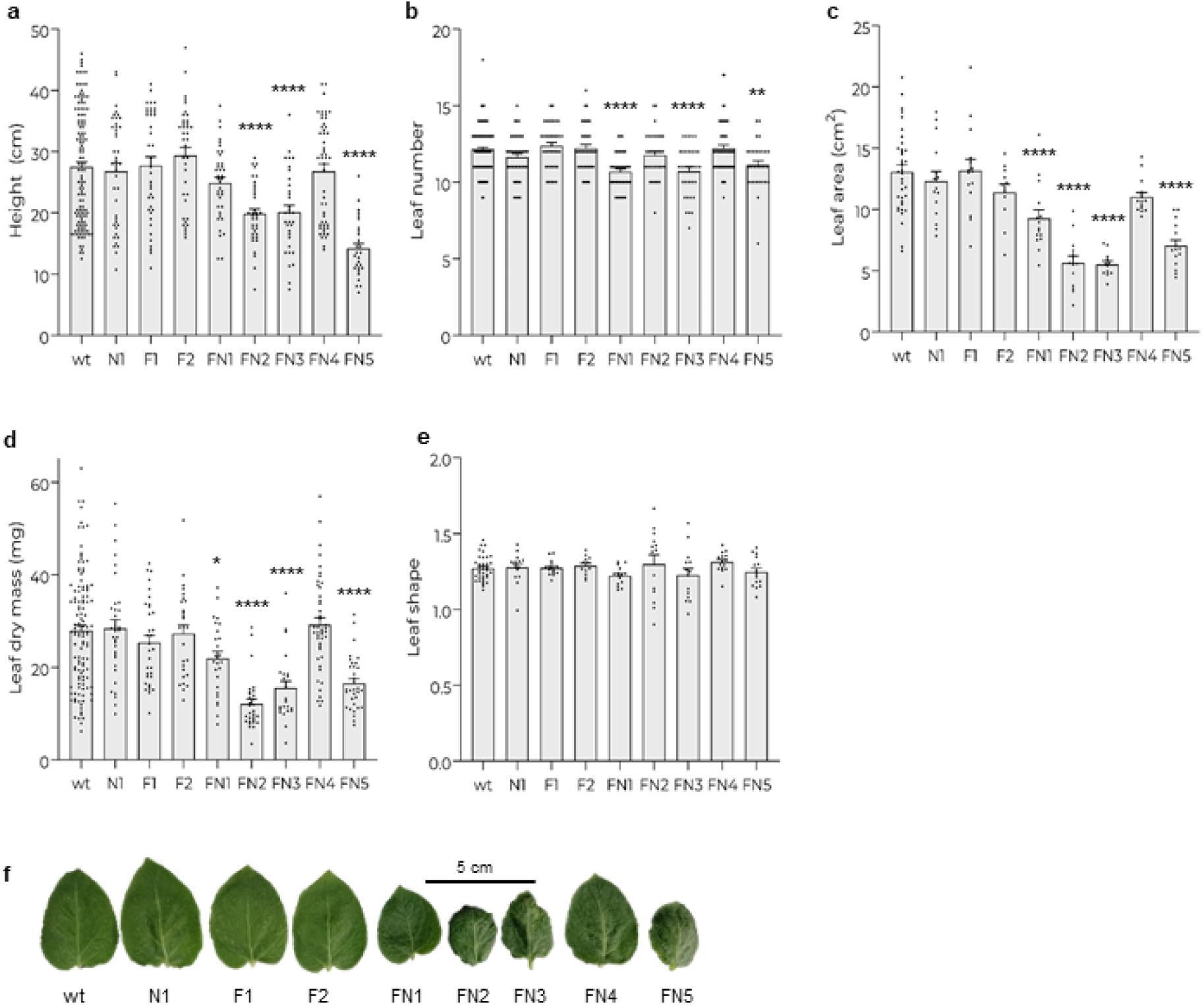
Vegetative growth. Wild type and transgenic plants were produced *ex vitro*. 4 weeks after soil transfer, vegetative growth parameters were determined. **a** Plant height. **b** Number of leaves per plant. **c**-**e** Area, dry mass, and shape (length width^-1^; **e**) of the apical leaflet of the second fully expanded leaf. **f** Representative image of the leaflets. **a**-**c** and **e** Data pooled from four distinct planting/harvest events, represented as individual measurements (dots: plants, or leaves from different plants) and mean ± s.e.m. **d** Data pooled from seven distinct planting/harvest events, represented as individual measurements (dots: leaves from different plants) and mean ± s.e.m. \**P* < 0.05; \*\**P* < 0.01; \*\*\*\**P* < 0.001, compared with the wt by one-way ANOVA with Bonferroni post-hoc test.

When evaluating tuberization parameters in *ex vitro* plants, FN1, FN2, FN3, and FN5 exhibited reduced yield and tuber weight compared to the wild type (Fig. 3a, b).

**Fig. 3.**
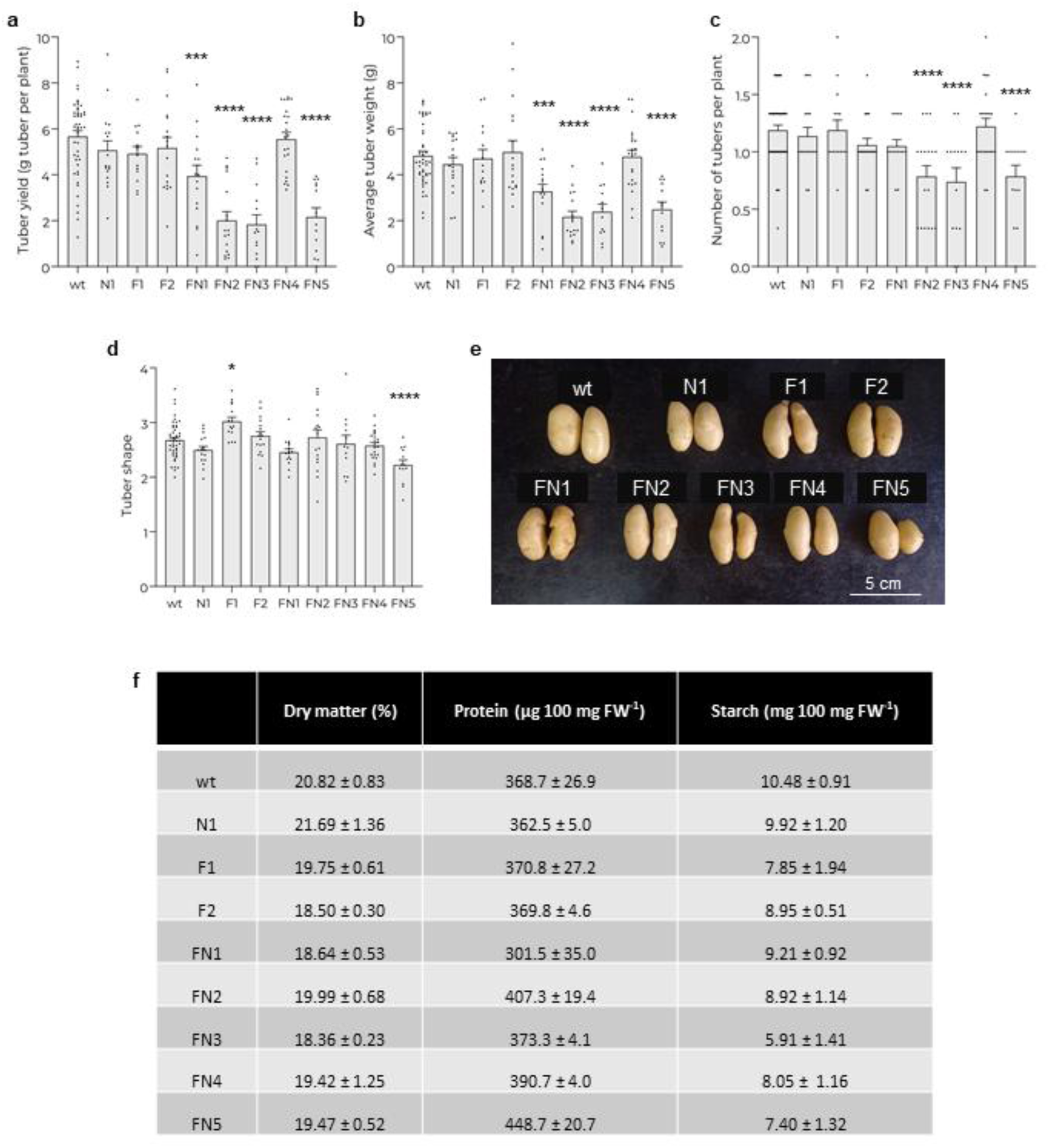
Yield and tuber characteristics. Wild type and transgenic plants were produced *ex vitro*. Tubers were harvested after plant senescence, and yield and tuber characteristics were determined. **a** Tuber yield. **b** Average tuber weight. **c** Number of tubers obtained per plant. **d** Tuber shape, defined as the ratio between the major and minor diameter. **e** Representative image of tubers. **f** Dry matter, protein, and starch content of tubers. **a**-**d**: Data pooled from four distinct planting/harvest events, represented as individual measurements (dots: plants, or average data of the tubers produced by one plant) and mean ± s.e.m. \**P* < 0.05; \*\**P* < 0.01; \*\*\**P* < 0.005; \*\*\*\**P* < 0.001, compared with the wt by one-way ANOVA with Bonferroni post-hoc test. **f**: Data are the mean ± s.e.m of at least three tubers from different plants from a single planting/harvest event; the differences between transgenic lines and wt were not significant for any of the parameters analyzed.

Furthermore, FN2, FN3, and FN5 showed a decrease in the number of tubers produced per plant (Fig. 3c). In contrast, these parameters remained unchanged in N1, F1, F2, and FN4 (Fig. 3a-c). Notably, tubers from F1 and FN5 plants displayed an altered shape (Fig. 3d, e). A similar trend in tuber yield was observed in plants developed from seed tubers (Supplementary Fig. S3a-c). However, under these conditions, more transgenic lines exhibited altered tuber shape (Supplementary Fig. S3d and e).

No significant differences were observed in dry matter, protein, or starch content between the wild type and any of the transgenic lines (Fig. 3f).

### Iron and zinc content

Iron and zinc content was assessed in tubers of wild type and transgenic plants obtained *ex vitro* (Fig. 4). Lines N1, F1, and F2 exhibited iron levels comparable to those of the wild type, whereas FN1, FN2, FN3, FN4 and FN5 showed increased iron concentrations. Similarly, the co-expressing lines displayed higher zinc levels compared to the wild type. The FN5 line exhibited stunted growth and a significantly reduced tuber yield at the onset of this study. Consequently, due to the limited availability of tuber material, this line was excluded from the zinc content analysis.

**Fig. 4.**
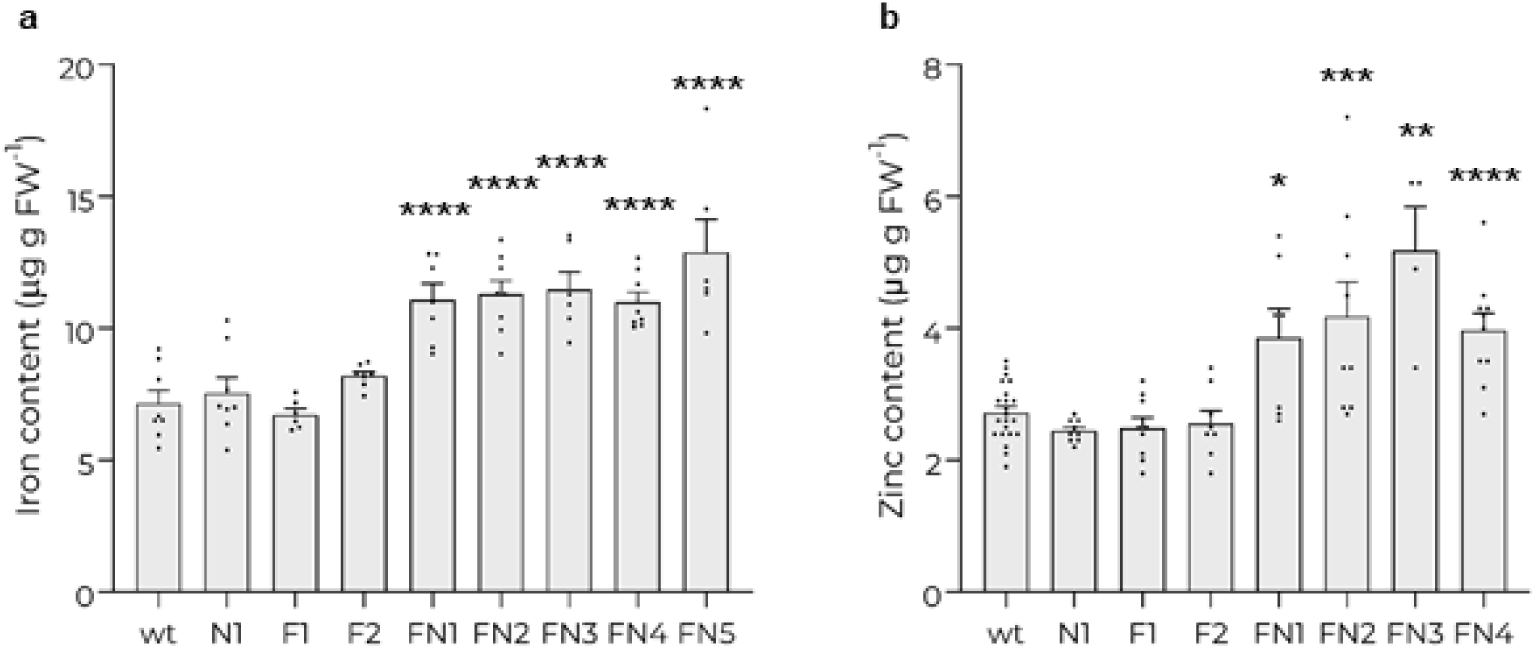
Iron and zinc content of tubers. Tubers of wild type and transgenic plants produced *ex vitro* were harvested after plant senescence. Iron content (**a**) was determined using the colorimetric method, and zinc content (**b**) was measured by atomic absorption spectrometry. **a** Data pooled from two independent planting/harvest events for wt, N1, F2, FN1, and FN4, and from one event for F1, FN2, FN3, and FN5. **b** Data pooled from three distinct planting/harvest events for wt, N1, F1, F2, FN2 and FN4, and from two events for FN1 and FN3. Data are presented as individual measurements (dots: tubers from different plants) and mean ± s.e.m. \**P* < 0.05; \*\**P* < 0.01; \*\*\**P* < 0.005; \*\*\*\**P* < 0.001, compared with the wt by one-way ANOVA with Bonferroni post-hoc test.

### Water loss, RWC and stomatal characteristics

Given the critical role of water balance in plant physiology and the observation that various co-expressing lines exhibit altered leaf morphology, the effect of *AtNAS1* and *PvFERRITIN* expression on water retention capacity was assessed. This evaluation involved measuring the rate of water loss and relative water content (RWC) in wild type and transgenic plants obtained *ex vitro*. As shown in Fig. 5a and 5b, leaves from transgenic lines N1, F1, FN2, FN3, and FN5 exhibited an increased rate of water loss compared to the wild type, whereas no significant effect was observed in F2, FN1, and FN4. Additionally, FN2, FN3 and FN5 also showed a decrease in RWC (Fig. 5c, d).

**Fig. 5.**
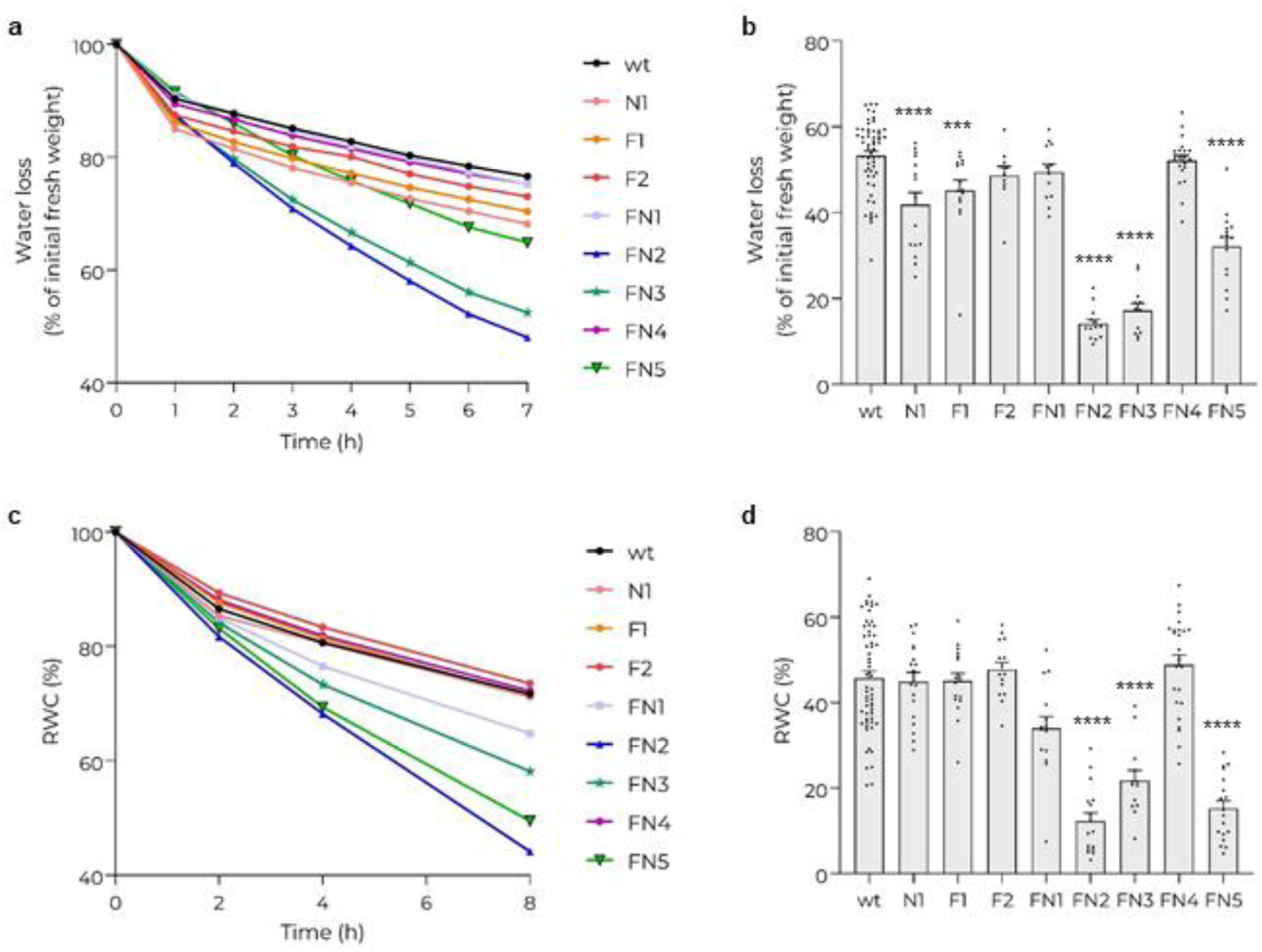
Water loss and RWC. **a** Water loss rate, determined in wild type and transgenic plants produced *ex vitro* 1-7 h after leaf detachment (statistical analysis is shown in Supplementary Fig. S4). **b** Water loss rate 24 h after leaf detachment; data pooled from three distinct planting/harvest events, represented as individual measurements (dots: leaves from different plants) and mean ± s.e.m; \*\*\**P* < 0.005; \*\*\*\**P* < 0.001, compared with the wt by one-way ANOVA with Bonferroni post-hoc test. **c** RWC, determined in wild type and transgenic plants produced *ex vitro* after 2, 4 and 8 h (statistical analysis is shown in Supplementary Fig. S5). **d** RWC after 24 h; data pooled from four distinct planting/harvest events, represented as individual measurements (dots: leaves from different plants) and mean ± s.e.m; \*\*\*\**P* < 0.001, compared with the wt by one-way ANOVA with Bonferroni post-hoc test.

To determine whether the reduced water retention capacity in the transgenic lines is due to an impaired stomatal function, the stomatal characteristics of the different lines were evaluated using plants obtained *ex vitro*. The lines FN2, FN3 and FN5 exhibited lower stomatal density and larger stomatal size compared to wild type plants, whereas these parameters were unaffected in N1, F1, F2, FN1, and FN4 (Fig. 6a, b, d). The stomatal index was reduced in FN2, FN3 and FN5, but this reduction was not statistically significant (Fig. 6c). When assessing plants developed from seed tubers, similar results were observed (Supplementary Fig. S6). However, under these conditions, the differences in stomatal index between the wild type and lines FN2, FN3, and FN5 were statistically significant.

**Fig. 6.**
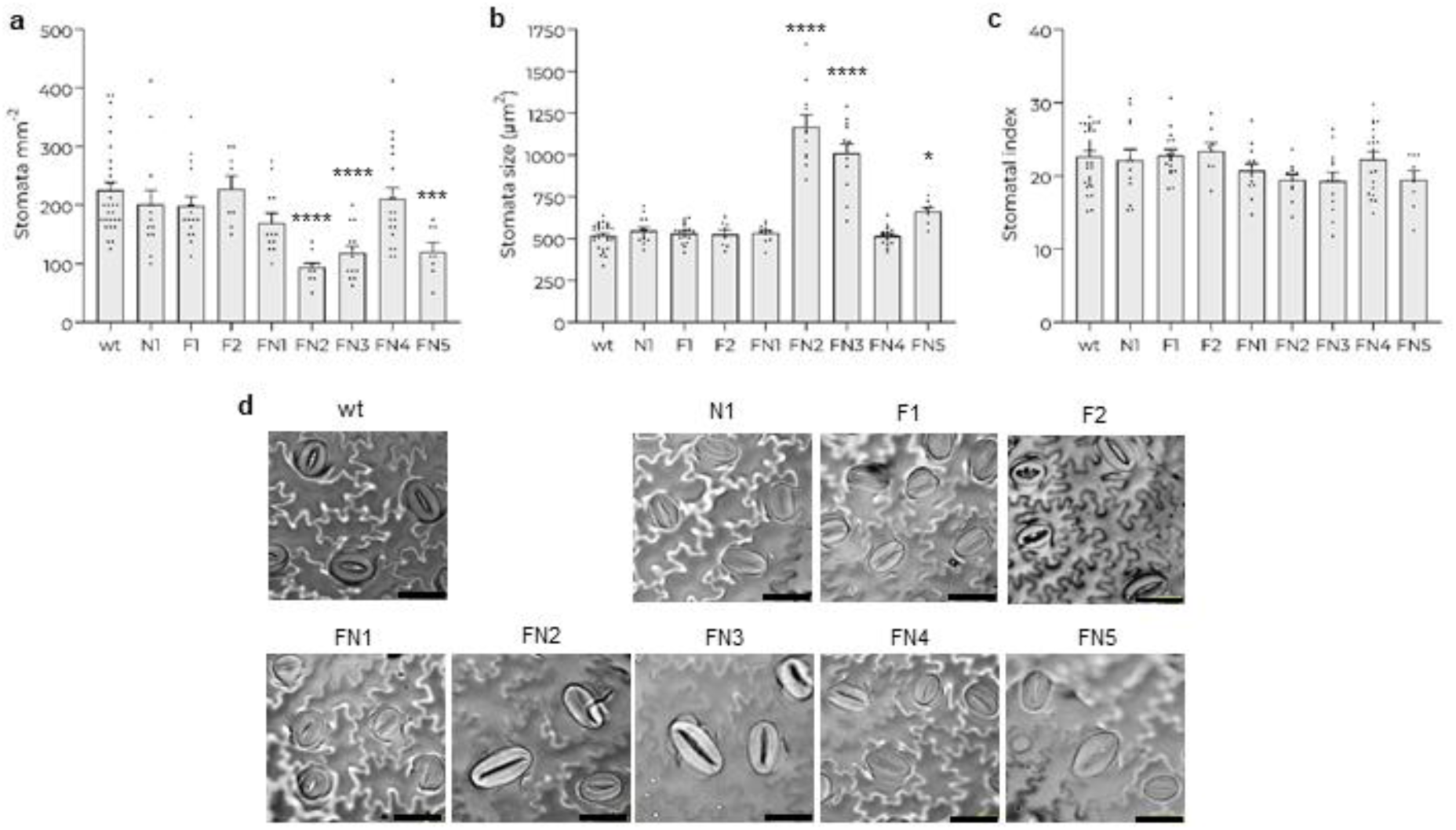
Stomatal characteristics. Stomatal density (**a**), stomatal size (**b**), and stomatal index (**c**) were determined in wild type and transgenic plants produced *ex vitro*. **d** Representative image of abaxial epidermis (scale bar: 40 μm). **a**-**c** Data pooled from three distinct planting/harvest events, represented as individual measurements (dots: leaves from different plants) and mean ± s.e.m; \**P* < 0.05; \*\*\**P* < 0.005; \*\*\*\**P* < 0.001, compared with the wt by one-way ANOVA with Bonferroni post-hoc test.

It has been observed that plants with a higher density of smaller stomata exhibit faster stomatal responses, whereas larger stomata present slower kinetics (Lawson and Matthews 2020). Therefore, the increased stomatal size and reduced stomatal density observed in lines FN2, FN3 and FN5 may contribute to their impaired ability to adjust stomatal movements, resulting in increased rates of water loss and decreased RWC.

### Photosynthetic pigments

When assessing the photosynthetic pigments chlorophyll and carotenoids in *ex vitro* plants, a significant decrease in total chlorophyll content was observed in lines FN1 and FN5 compared to the wild type, due to reductions in chlorophyll a and b (Fig. a-c). FN2, and FN3 showed a same trend, but the differences from the wild type were not statistically significant. In contrast, chlorophyll content remained unaffected in lines N1, F1, F2, and FN4. No differences were observed in the chlorophyll a to chlorophyll b ratio among the transgenic lines and the wild type (Fig. 7d). The total carotenoid content remained unchanged in all transgenic lines (Fig. 7e). Similar results were obtained for plants grown from seed tubers (Supplementary Fig. S7). Under these conditions, lines FN1, FN3, and FN5 exhibited a statistically significant reduction in total chlorophyll content due to decreases in chlorophyll a and b. Additionally, FN5 exhibited a reduction in total carotenoid content.

**Fig. 7.**
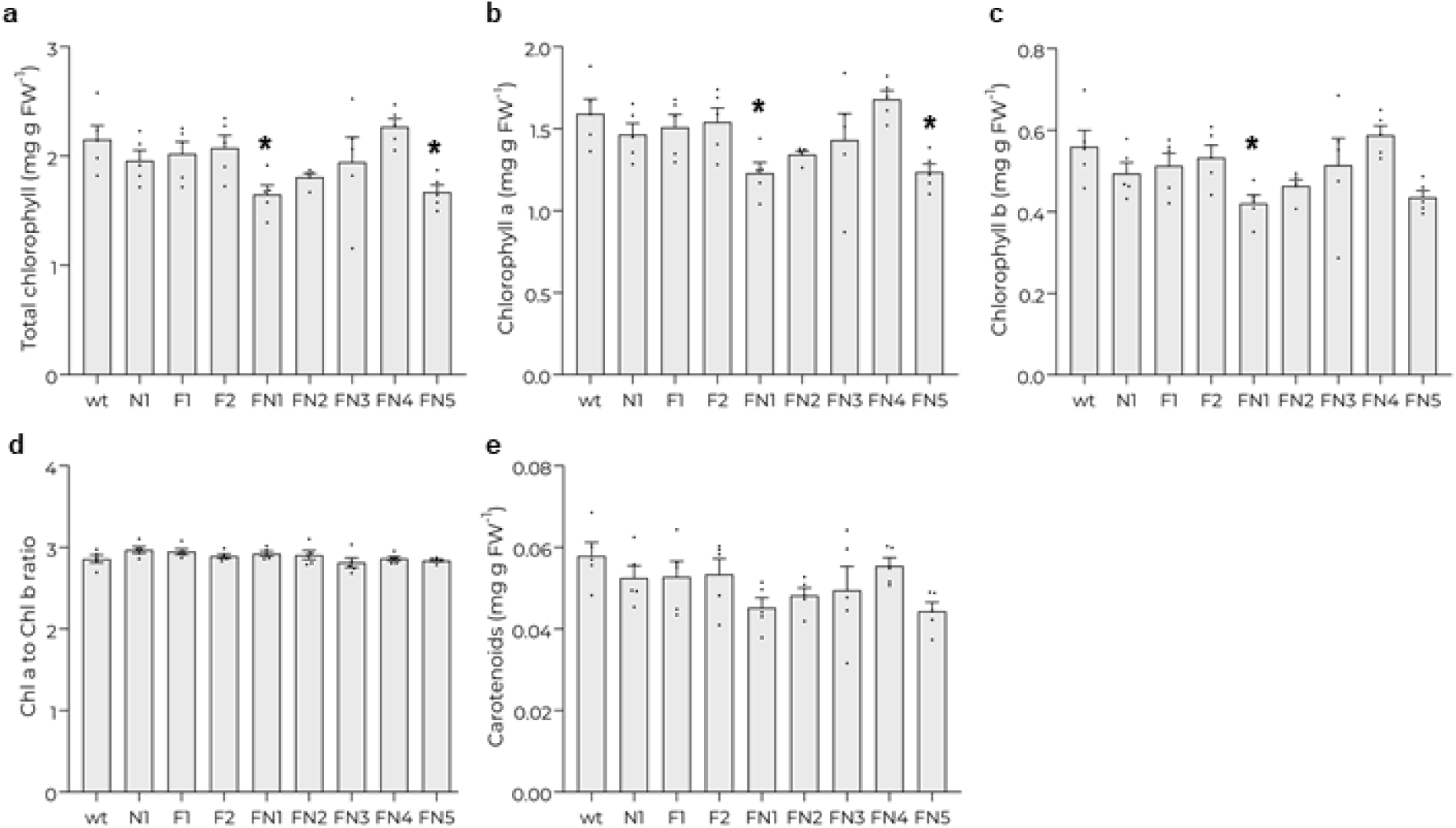
Photosynthetic pigments content. Total chlorophyll, chlorophyll a, and chlorophyll b content (**a**-**c**); chlorophyll a to chlorophyll b ratio (**d**), and total carotenoids content (**e**), determined in leaves from wild type and transgenic plants produced *ex vitro*. Data are the mean ± s.e.m of individual measurements (dots: leaves from different plants) from a single planting/harvest event. \**P* < 0.05, compared with the wt by one-way ANOVA with Bonferroni post-hoc test.

### Assessment of line FN4 grown in soil

The assessment of morphological and physiological parameters, along with iron and zinc content, was conducted on plants grown either *ex vitro* or from seed tubers in Klassman TS 085 substrate. This substrate is composed of Sphagnum peat moss, fertilizer, and trace elements, providing a stable composition that is essential for ensuring experimental reproducibility. From this evaluation, we determined that the FN4 line exhibits increased iron and zinc content in tubers, without compromising yield or affecting any of the assessed morphological or physiological parameters (Fig. 2-7).

We evaluated the performance of the FN4 line using plants grown from seed tubers in a soil and organic compost mixture, simulating conditions more comparable to field cultivation. Under these growth conditions, FN4 plants showed no significant differences compared to wild type plants in terms of plant height (Fig. 8a), number of leaves per plant (Fig. 8b), tuber yield (Fig. 8d), tuber weight (Fig. 8e), number of tubers per plant (Fig. 8f), or tuber shape (Fig. 8g). However, FN4 plants exhibited an increased chlorophyll concentration compared to the wild type (Fig. 8c). The iron and zinc contents were significantly higher in tubers from FN4 plants than in those from wild-type plants (Fig. 8h, i), with increases of 111.4% for iron and 79.1% for zinc.

**Fig. 8.**
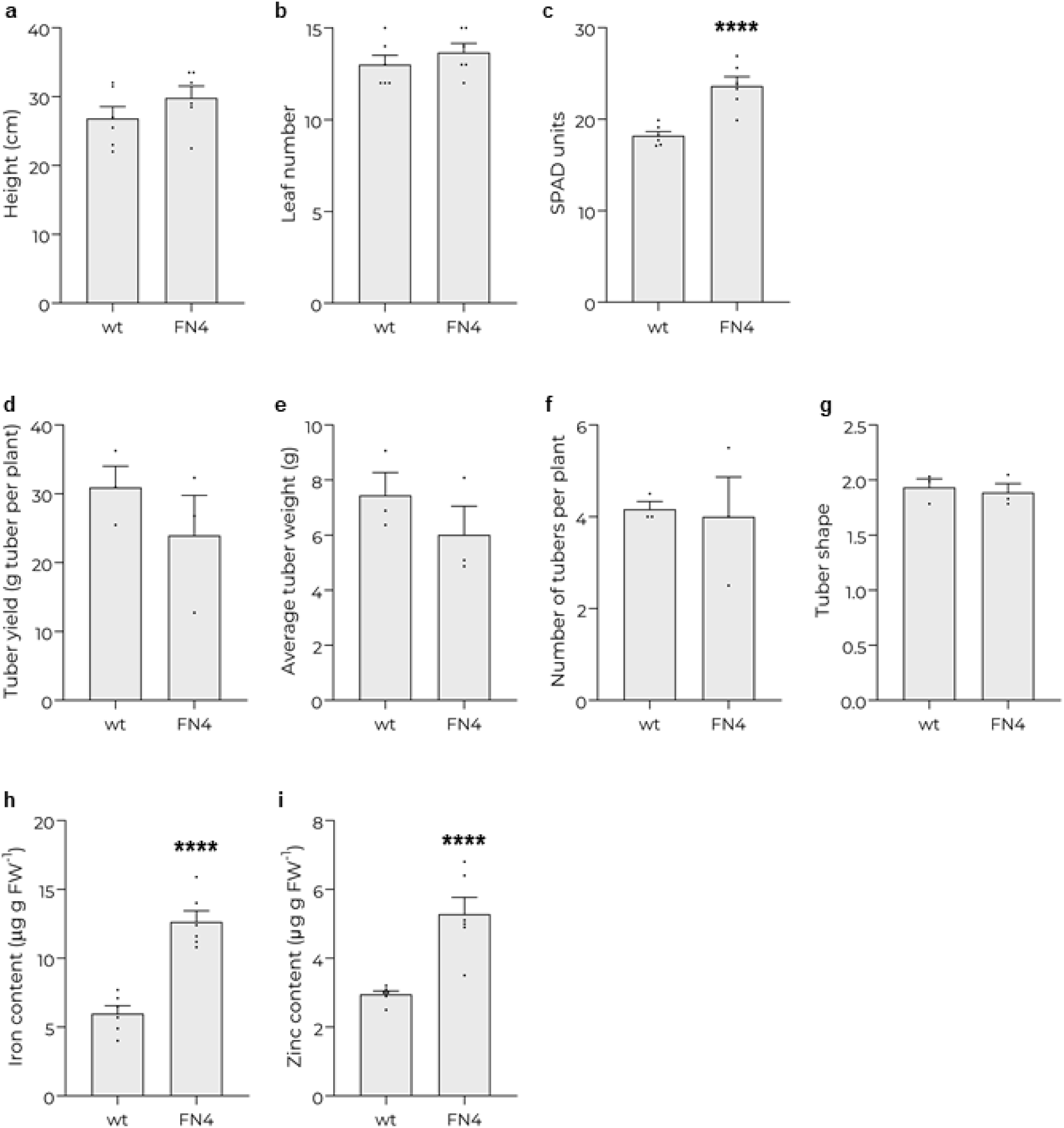
Assessment of line FN4 grown in soil. Wild type and FN4 plants developed from seed tubers were cultivated in pots filled with soil-compost mixture. 4 weeks after planting, the plant height (**a**), the number of leaves per plant (**b**), and the chlorophyll content (SPAD units; **c**) were measured. Data are the mean ± s.e.m of individual measurements (dots: plants, or leaves from different plants) from a single planting/harvest event. Tubers were harvested after plant senescence, and the tuber yield (**d**), the average tuber weight (**e**), the number of tubers obtained per plant (**f**), and the tuber shape, defined as the ratio between the major and minor diameter (**g**) were determined. Data are the mean ± s.e.m of individual measurements (dots: plants, or average data of the tubers produced by one plant) from a single planting/harvest event. **h** and **i** Iron and zinc content of tubers, measured by atomic absorption spectrometry. Data are the mean ± s.e.m of 6 tubers from different plants from a single planting/harvest event. \*\*\*\**P* < 0.001, compared with the wt by Student‘s *t* test.

### Effects of soil iron concentration on growth, yield and tuber iron content

We evaluated the growth, yield, and iron accumulation in tubers of wild-type and FN4 plants using *ex vitro* plants cultivated in Klassman TS 085 substrate, with or without the addition of varying amounts of Fe-EDDHA. The addition of 5 g of Fe-EDDHA did not significantly affect plant height (Fig. 9a), number of leaves (Fig. 9b), or yield (Fig. 9d) in either wild type or FN4 plants. In contrast, the addition of 10 g of Fe-EDDHA negatively impacted growth in both wild type and FN4 plants (Fig. 9a), but only reduced yield in the wild type (Fig. 9d). Chlorophyll content increased in response to Fe-EDDHA addition in the leaves of both wild-type and FN4 plants (Fig. 9c).

**Fig. 9.**
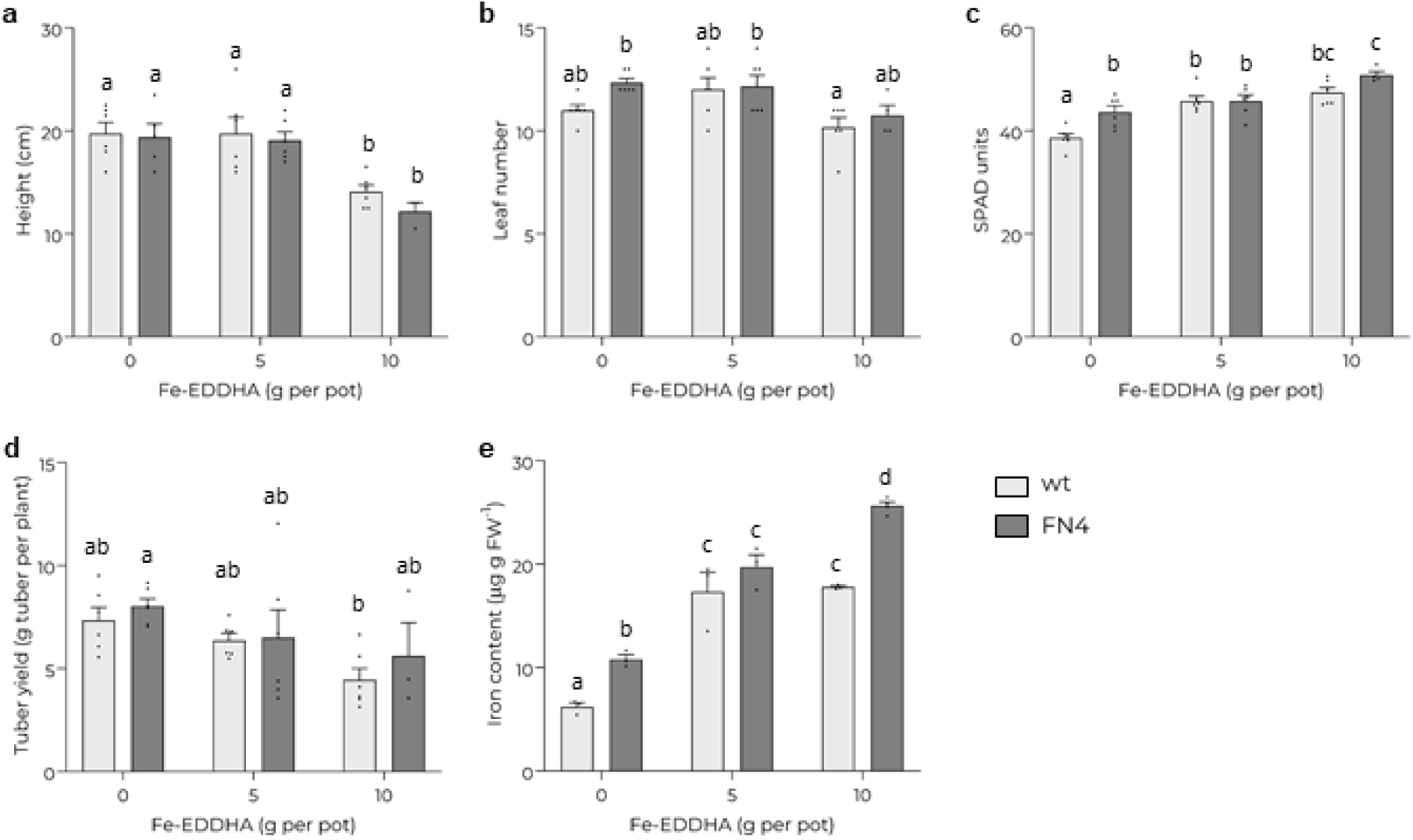
Effects of soil iron concentration on growth, yield and tuber iron content. Wild type and FN4 plants produced *ex vitro* were cultivated in Klassman TS 085 substrate, with or without the addition of the indicated amounts of Fe-EDDHA. 4 weeks after soil transfer, the plant height (**a**), number of leaves per plant (**b**), and SPAD units (**c**) were determined. Tubers were harvested after plant senescence; the tuber yield was determined (**d**), and tuber iron content was measured by the colorimetric method (**e**). Data are the mean ± s.e.m of individual measurements (dots: plants, average data of the tubers produced by one plant, or tubers from different plants) from a single planting/harvest event. Data were analyzed by two-way ANOVA, followed by Tukeýs post-hoc test. Different letters in the bar graphs indicate statistically significant differences.

Interestingly, tuber iron content significantly increased in both wild type and FN4 plants with the addition of 5 g of Fe-EDDHA. However, while the addition of 10 g of Fe-EDDHA further enhanced iron content in FN4 tubers, no such increase was observed in wild type tubers (Fig. 9e). These results indicate that line FN4 has a greater capacity to accumulate iron in tubers compared to the wild type, with iron accumulation in the wild type saturating at lower soil iron concentrations.

## DISCUSSION

The aim of this study was to develop biofortified varieties of the potato cultivar Spunta with enhanced iron and zinc content in the tubers. To achieve this, transgenic potato plants were generated expressing *AtNAS1*, *PvFERRITIN*, or both genes (Fig. 1). Remarkably, significant increases in iron and zinc concentrations were observed exclusively in lines co-expressing both genes (Fig. 4). These findings reveal that simultaneous expression of *AtNAS1* and *PvFERRITIN* is required to achieve substantial enhancements in iron and zinc levels in Spunta potato tubers.

A recent study demonstrated that overexpression of *AtNAS1* alone in the potato variety Désirée increased tuber iron content by 2.4-fold (Zha et al. 2022). Using a similar approach, we generated one Spunta line (N1), which showed no differences in tuber iron or zinc content compared to the wild type (Fig. 4). However, to draw robust conclusions about the effectiveness of this strategy, it would have been necessary to obtain additional Spunta lines overexpressing *AtNAS1* alone.

The phenotypic evaluation of the transgenic lines revealed that co-expression of *AtNAS1* and *PvFERRITIN* affects both vegetative growth and tuber yield (Fig. 2, Fig. 3a-c). Among the lines expressing both genes, FN4 exhibited the least adverse effects, with growth and tuberization parameters closely resembling those of the wild type. Furthermore, our findings indicate that *AtNAS1* and *PvFERRITIN* expression in potato impairs the plant’s water retention capacity possibly by altering stomatal density and size (Fig. 5, Fig. 6). The reduction in stomatal density, accompanied by an increase in stomatal size in lines FN2, FN3, and FN5, along with smaller leaf sizes and reduced leaf numbers, suggests that the introduction of these genes impacts multiple aspects of leaf development. Additionally, co-expression of *AtNAS1* and *PvFERRITIN* influences photosynthetic pigment concentration (Fig. 7). Notably, FN4 showed no significant changes in water loss rate, RWC, or stomatal characteristics, while exhibiting increased chlorophyll levels (Fig. 8c; Fig. 9c).

Tubers of the FN4 line exhibited a greater capacity to accumulate iron than wild type tubers under conditions of increased soil iron availability, without affecting plant yield, even when vegetative growth rates were reduced due to elevated soil iron levels (Fig. 9). This outcome highlights the effectiveness of introducing the *PvFERRITIN* gene under the control of a tuber-specific promoter, in combination with the *AtNAS1* gene, to enhance iron uptake, transport, and targeted accumulation in the tubers.

Our findings demonstrate that the co-expression of *AtNAS1* under a constitutive promoter and *PvFERRITIN* under a tuber-specific promoter is a promising strategy for improving iron and zinc content in the tubers of the Spunta potato variety. However, given the potential effects of these genes on growth, development, and yield, it is crucial to evaluate multiple transformation events to select one that offers effective biofortification with minimal negative impacts. In this study, among five co-expressing events, we identified FN4, which exhibits a 2.1-and 1.8-fold increase in tuber iron and zinc content, respectively, without compromising yield (Fig. 8). Additionally, our findings indicate that iron accumulation in tubers is influenced by the substrate in which the plants are cultivated (Fig. 4; Fig. 8; Fig. 9). Therefore, when assessing strategies for iron biofortification in potato, it is important to test different substrates.

The biofortified FN4 potato variety could be an important dietary source of iron. For women aged 19 to 50, the recommended dietary allowance (RDA) for iron is 18.0 mg/day, assuming an absorption rate of 18% (Institute of Medicine 2001). Considering an absorption rate of 28.4%, as found in yellow-fleshed potatoes (Jongstra et al. 2020), a 300 g serving of FN4 tubers (equivalent to the average weight of a Spunta tuber) would provide 33.3% of the daily iron requirement. Additionally, the same portion would fulfill 19.8% of the daily zinc RDA for women. This variety holds significant potential to address iron deficiency anemia and zinc deficiency, improving the health of populations in both rural and urban areas affected by poverty. Field testing is necessary to confirm the agronomic resilience and nutritional benefits of this high-iron/high-zinc potato variety.

## Supplementary Information

The online version contains supplementary material.

## Acknowledgements

We thank Navreet K. Bhullar for his kind gift of the IINF vector.

## Author contributions

DAC and MNMG conceived and designed the research. JIC, MZ, VRS, and MF conducted the experiments and analyzed the data. DAC wrote the manuscript. All authors read and approved the final version.

## Data Availability

The data that support this study are available in the article and accompanying online supplementary material.

## Declarations

