## Supplemental files for "Iron and zinc biofortification in potato through the introduction of *NICOTIANAMINE SYNTHASE* and *FERRITIN* genes"

**Table S1.** Primers used in this study.

| | Primer sequence |
|---|---|
| AtNAS1 forward | ACTCGTCTGATCTCAAAGGC |
| AtNAS1 reverse | CTCGATGGCACTAAACTCCTCG |
| PvFERRITIN forward | AGATCGCAACAAAGACCCTC |
| PvFERRITIN reverse | CATATGATACTCGCTACCAC |
| NPTII forward | ATGATTGAAGAAGATGGATTG |
| NPTII reverse | GAAGAACTCGTCAAGAAGGCG |
| EF1-α forward | GTATGGTTGTGACCTTTGG |
| EF1-α reverse | CAACATTCTTGACAACAC |

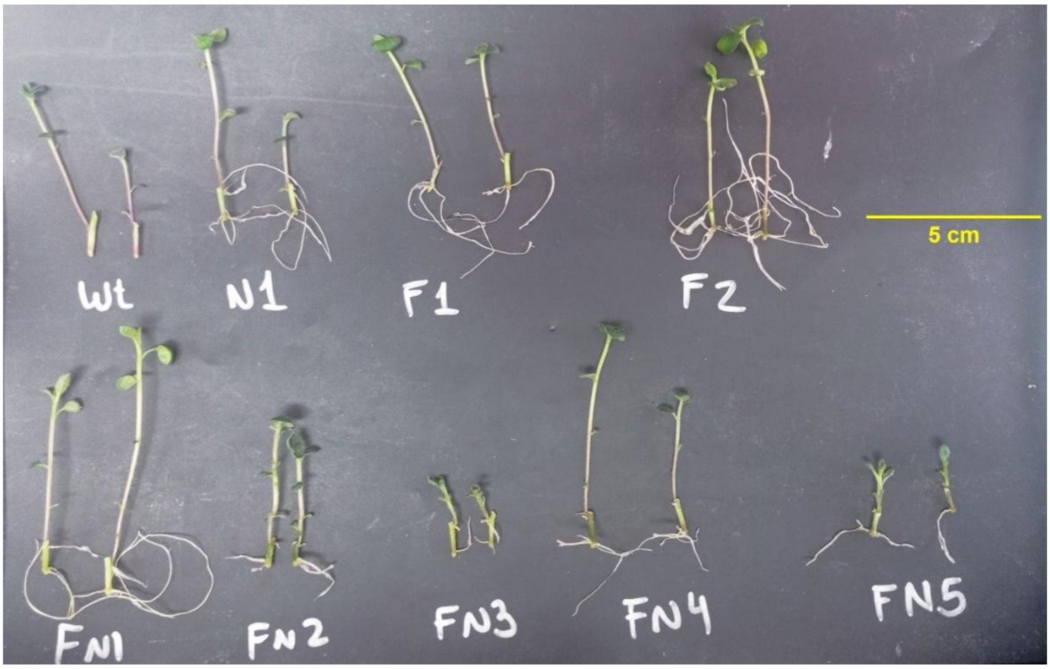


**Fig. S1.** Root development in selective media. Single-node cuttings from wild type (wt) and transgenic plants (N1, F1-2, FN1-5) were cultured on MS medium with 20 g/L sucrose, solidified with 0.7% (w/v) agar, containing 50 mg/L kanamycin, in a growth chamber under a 16-h light photoperiod (4000 lx light intensity) at 22 °C. After 2 weeks, plants were photographed to evaluate root development. A representative image is shown.

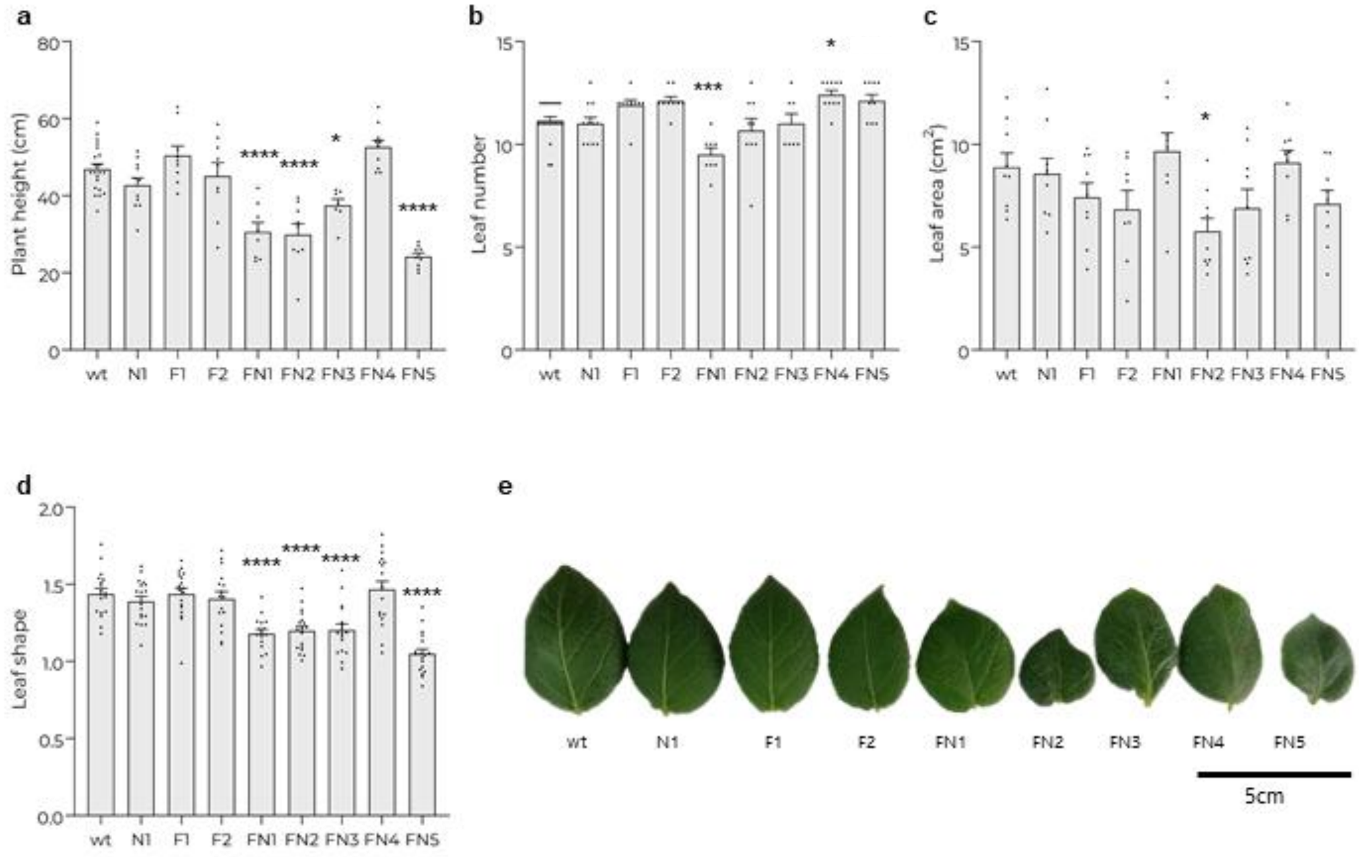


**Fig. S2.** Vegetative growth. Wild-type and transgenic plants were developed from seed tubers. 4 weeks after planting vegetative growth parameters were determined. **a** Plant height. **b** Number of leaves per plant. **c** and **d** Area and shape (length width$^{-1}$) of the apical leaflet of the second fully expanded leaf. **e** Representative image of the leaflets. Data are the mean ± s.e.m of individual measurements (dots: plants, or leaves from different plants) from a single planting/harvest event. *$P$ < 0.05; ***$P$ < 0.005; ****$P$ < 0.001, compared with the wt by one-way ANOVA with Bonferroni post-hoc test.

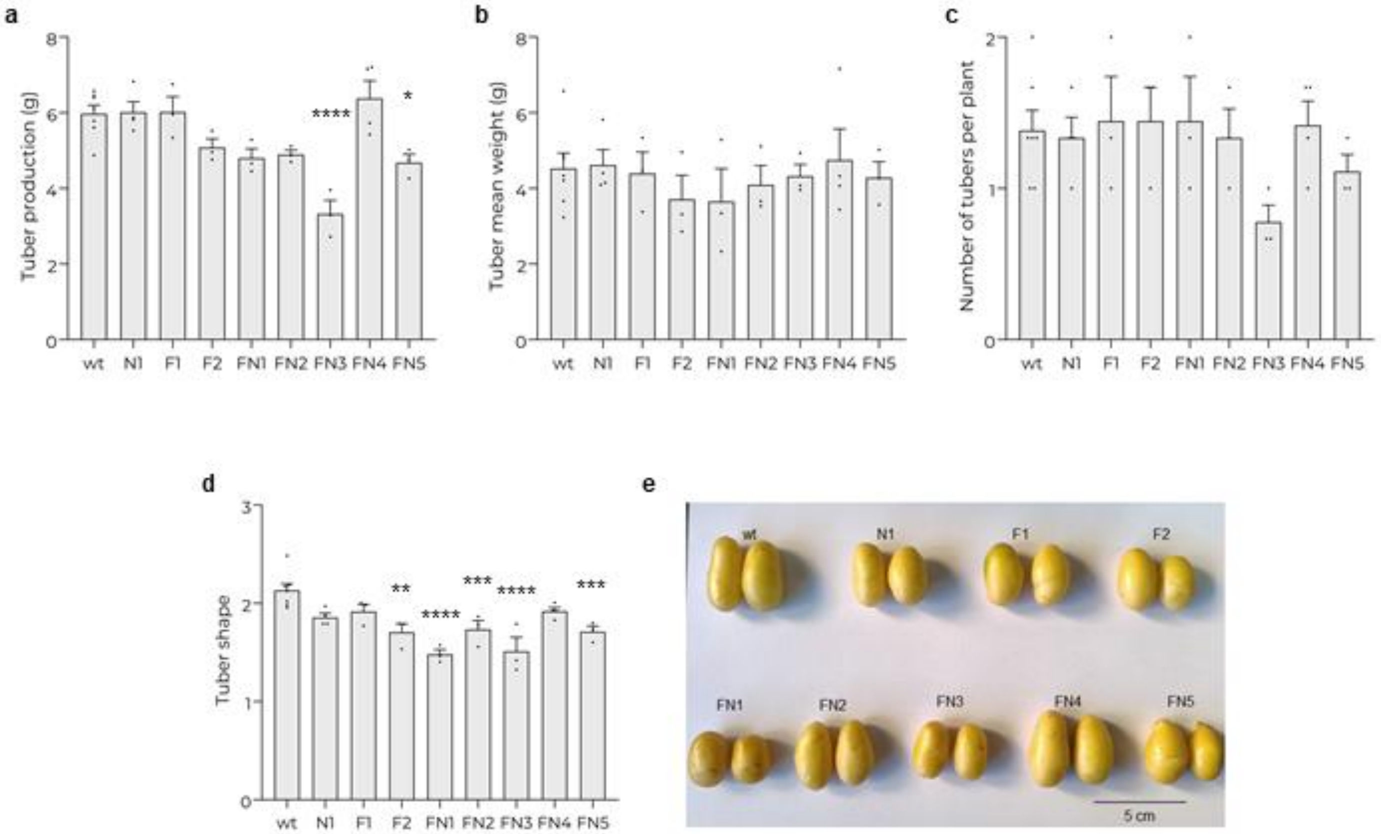


**Fig. S3.** Yield and tuber characteristics. Wild type and transgenic plants were developed from seed tubers. Tubers were harvested after plant senescence, and yield and tuber characteristics were determined. **a** Tuber yield per plant. **b** Average tuber weight. **c** Number of tubers obtained per plant. **d** Tuber shape, defined as the ratio between the major and minor diameter. **e** Representative image of tubers. Data are the mean ± s.e.m of individual measurements (dots: plants, or average data of the tubers produced by one plant) from a single planting/harvest event. *$P < 0.05$; **$P < 0.01$; ***$P < 0.005$; ****$P < 0.001$, compared with the wt by one-way ANOVA with Bonferroni post-hoc test.

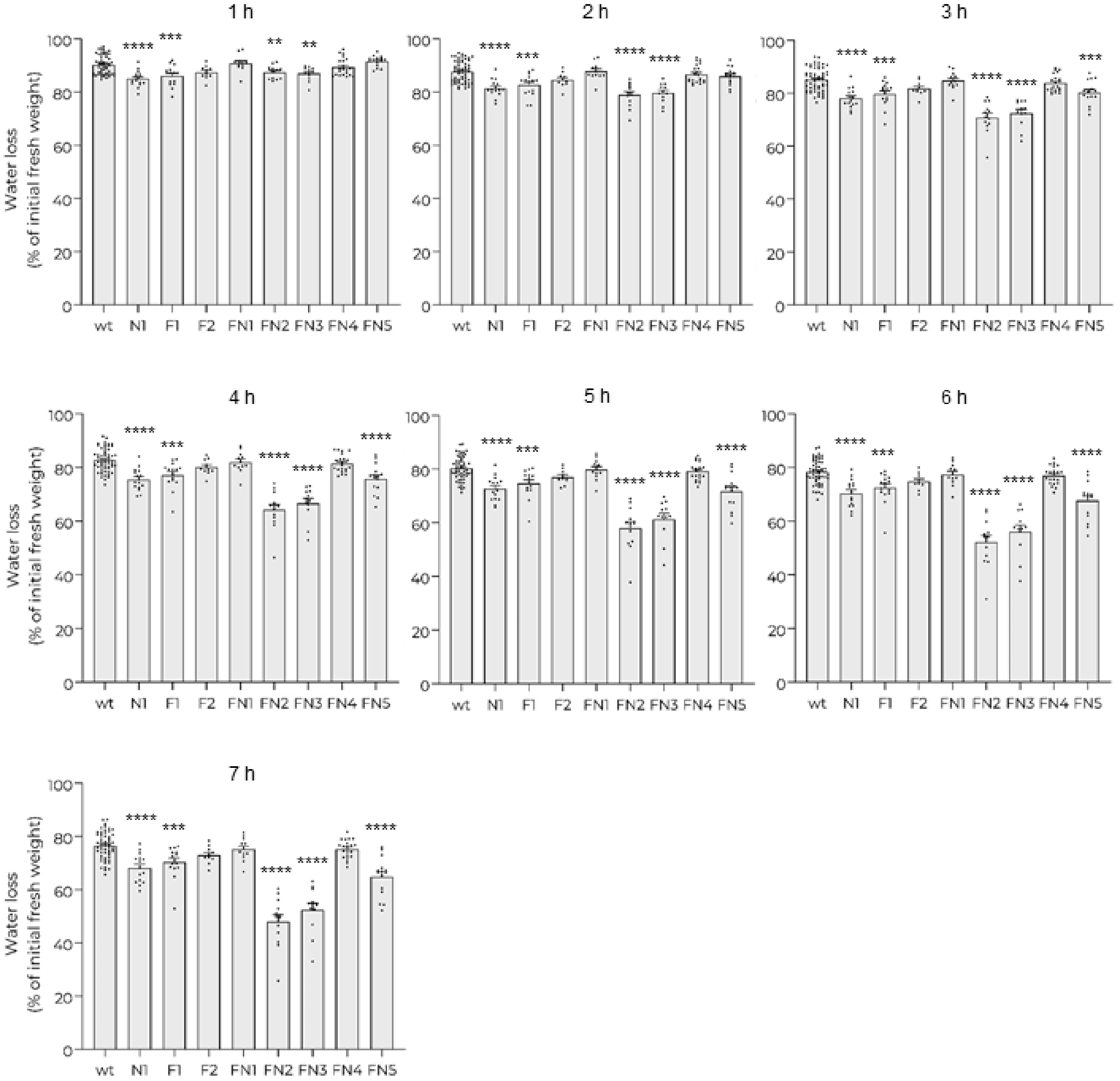


**Fig. S4.** Statistical analysis of Fig. 5a. Data pooled from three distinct planting/harvest events, represented as individual measurements (dots: leaves from different plants) and mean ± s.e.m; **$P < 0.01$; ***$P < 0.005$; ****$P < 0.001$, compared with the wt by one-way ANOVA with Bonferroni post-hoc test.

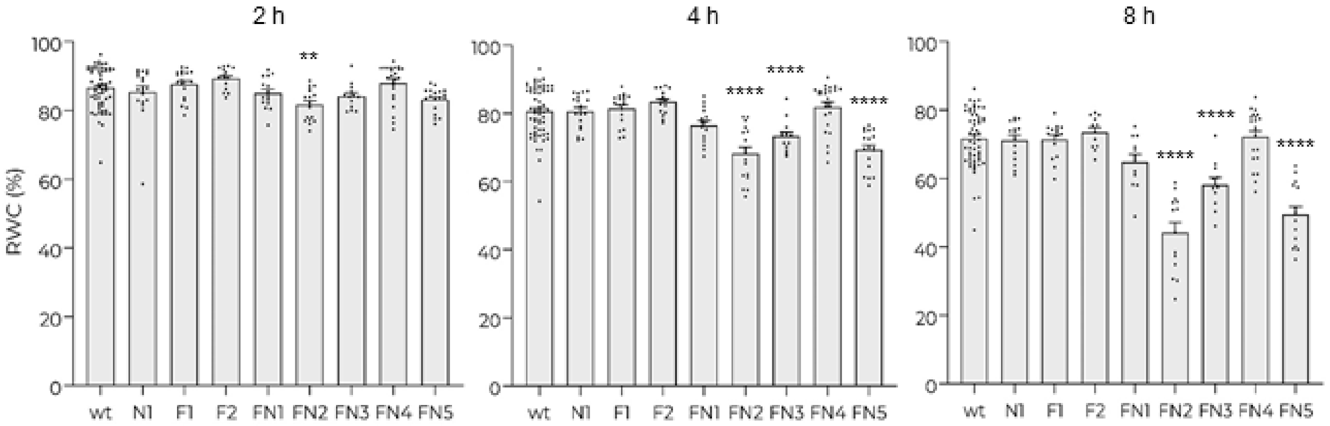


**Fig. S5.** Statistical analysis of Fig. 5b. Pooled data from four distinct planting/harvest events, represented as individual measurements (dots: leaves from different plants) and mean ± s.e.m; **$P$ < 0.01; ****$P$ < 0.001, compared with the wt by one-way ANOVA with Bonferroni post-hoc test.

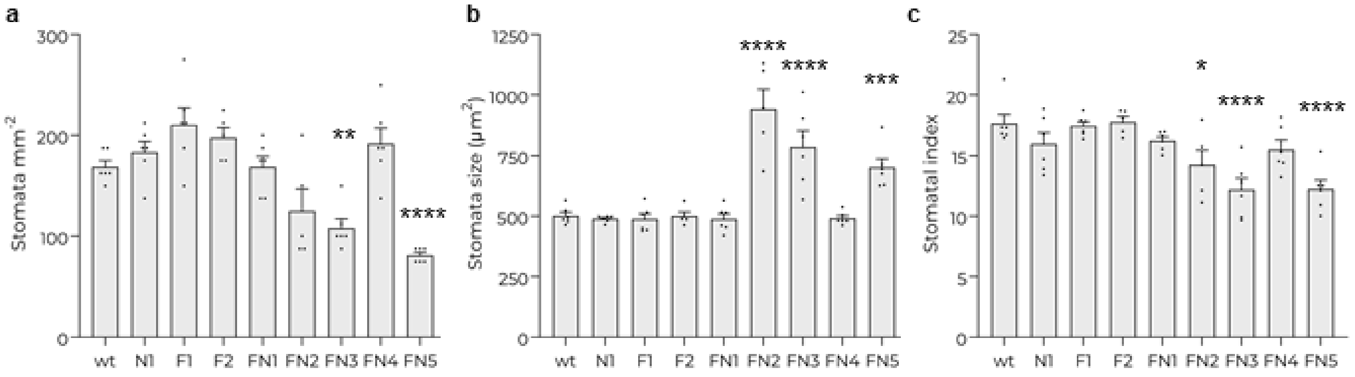


**Fig. S6.** Stomatal characteristics. Stomatal density (**a**), stomatal size (**b**), and stomatal index (**c**) were determined in wild type and transgenic plants developed from seed tubers. Data are the mean ± s.e.m of individual measurements (dots: leaves from different plants) from a single planting/harvest event. *$P$ < 0.05; **$P$ < 0.01; ***$P$ < 0.005; ****$P$ < 0.001, compared with the wt by one-way ANOVA with Bonferroni post-hoc test.

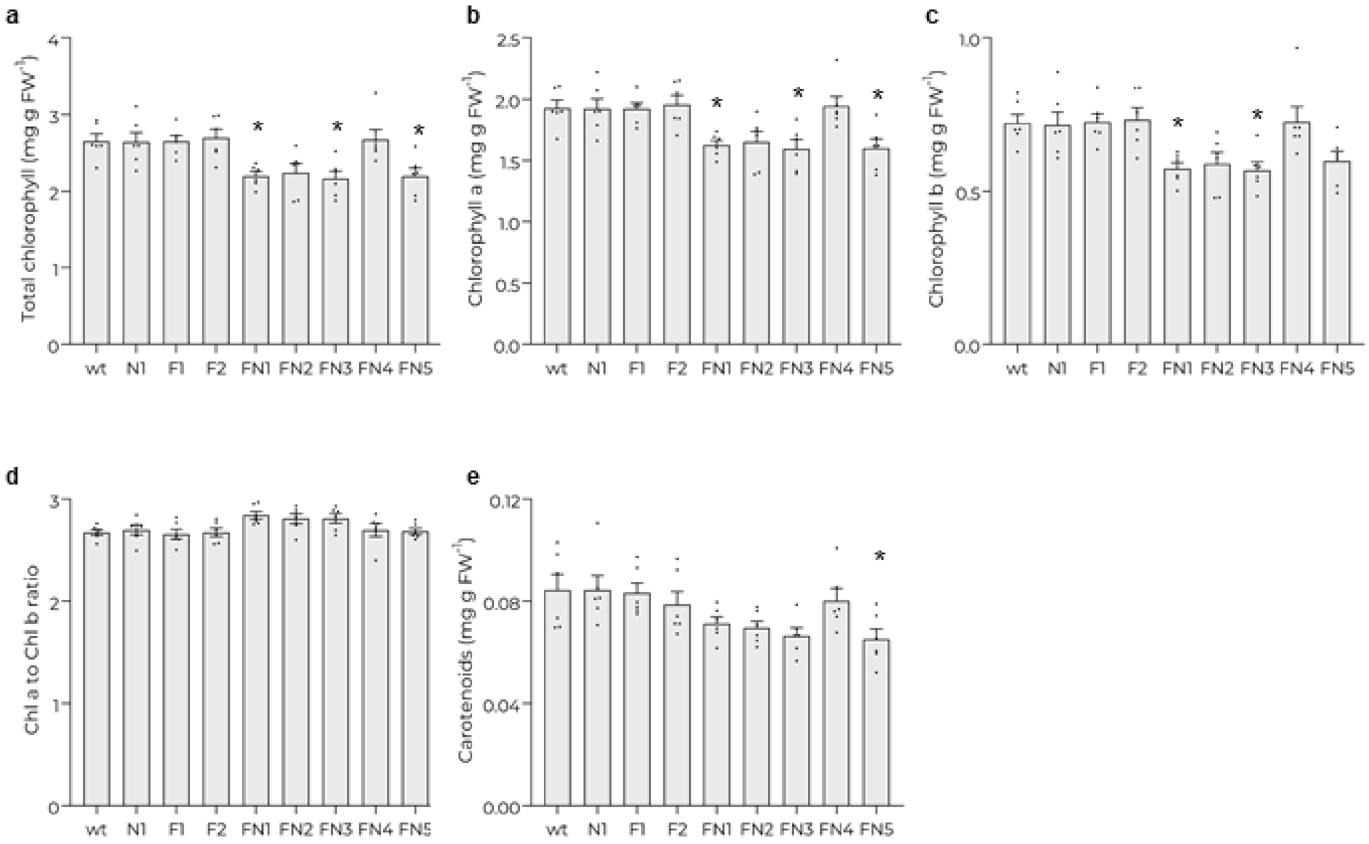


**Fig. S7.** Photosynthetic pigments content. Total chlorophyll, chlorophyll a, and chlorophyll b content (**a**-**c**); chlorophyll a/chlorophyll b ratio (**d**), and total carotenoids content (**e**), determined in leaves from wild type and transgenic plants developed from seed tubers. Data are the mean ± s.e.m of individual measurements (dots: leaves from different plants) from a single planting/harvest event. *$P < 0.05$, compared with the wt by one-way ANOVA with Bonferroni post-hoc test.
